# Developing heterospecific Sterile Insect Technique for pest control: insights from the spotted wing fly *Drosophila suzukii*

**DOI:** 10.1101/2024.09.05.611447

**Authors:** Flavia Cerasti, Massimo Cristofaro, Valentina Mastrantonio, Jessica Scifo, Adriano Verna, Daniele Canestrelli, Daniele Porretta

**Affiliations:** Department of Environmental Biology, Sapienza University of Rome, Rome, Italy; Biotechnology and Biological Control Agency (BBCA), Rome, Italy; FSN-FISS-SNI Laboratory, Italian National Agency for New Technologies, Energy and Sustainable Economic Development (ENEA), Rome, Italy; Department of Ecology and Biology, Tuscia University, Viterbo, Italy

**Keywords:** Reproductive interference, pest control, Sterile Insect Technique, *Drosophila suzukii*, *Drosophila melanogaster*

## Abstract

**BACKGROUND:** Reproductive interference (i.e., sexual interaction between males of one species and females of another species that reduce the fitness of one or both the interacting individuals) is an important species interaction significantly affecting population dynamics and persistence. However, its exploitation in pest control remains overlooked. Here, we investigated the possible integration of reproductive interference into the Sterile Insect Technique (SIT) to develop a heterospecific SIT (h-SIT). Under this approach, contrary to the classic SIT, sterile heterospecific males from closely related, non-pest species are released to compete with the pest population for mates. At this aim, we focused on the invasive pest species *Drosophila suzukii* and used *D. melanogaster* as the control species. First, we investigated the effect of irradiation on *D. melanogaster* sterility and longevity. Then, we tested the mating performance of irradiated males and their ability to reduce the *D. suzukii* fitness.

**RESULTS:** We found by microcosm experiments that: i) irradiation induced high levels of *D. melanogaster* male sterility without reducing longevity; ii) irradiated *D. melanogaster* males court *D. suzukii* females as much as *D. suzukii* males and they couple, mate and fecund heterospecific females; iii) irradiated *D. melanogaster* males significantly reduce the offspring of *D. suzukii* females under two different species ratios.

**CONCLUSIONS:** Our results provide the first foundation to develop a heterospecific Sterile Insect Technique against *D. suzukii* and fuel to test this approach against other groups of pest species.

## 1. Introduction

Species interactions, such as predation, parasitism, or competition, can significantly affect the population dynamics of the interacting species, affecting their abundance and persistence [1]. These interactions have been used for a long time to control pest species in agriculture and are the foundation of modern biological control approaches [2].

Along with the above species interactions, reproductive interference is now recognized as a major ecological process affecting population dynamics and species persistence [3–7]. Reproductive interference (or satyrisation in animals) consists of any sexual interaction between species that reduce the fitness of one or both interacting individuals [8]. It occurs due to incomplete mating barriers between species and can occur at any stage of mate acquisition, from courtship to mating and hybridization [3]. Reproductive interference has been documented under laboratory and field conditions in a wide variety of sexually reproducing taxa [5,6]. Theoretical and empirical studies showed that reproductive interference, as competition, is density-dependent and can result in population or species exclusion [4,9].

The first attempts to exploit reproductive interference for pest control date back to the first mid of the XX century. During the 1930s – 40s, in pioneering works, F. L. Vanderplank and colleagues applied reproductive interference to suppress tsetse fly, *Glossina swynnertoni* Austen (reviewed in [10]). Laboratory experiments showed that offspring produced by cross-mating between *Glossina swynnertoni* (the vector species) and *G. morsitans* Westwood (a nonvector species) had low fertility, as all hybrid males and some hybrid females were sterile. Then, a large field experiment was carried out in the Shinyanga area, Tanzania, by releasing fertile *G. morsitans* pupae over seven months. After the mass release, the density of *G. swynnertoni* was drastically reduced demonstrating the potential of hybrid sterility and sexual interference to suppress tsetse flies [9,10].

Although the experience with tsetse flies showed the potential effectiveness of reproductive interference for controlling a pest species, it did not lead to common exploitation of this approach. A major concern about heterospecific sterility was that pre-mating isolation mechanisms could constrain mating between released heterospecific males and wild females, leading to failure to control the target species in the field [11,12]. A further concern has been the risk of introducing non-native pest species. The mass release of fertile individuals of a closely related species that is non-native or a pest itself, at least potentially, would lead to replacing one pest species with another. These concerns and the success in the 1950s of Knipling and his team in eradicating the screwworm *Cochliomyia hominivorax* (Coquerel) using ionizing radiation [13], has led to the spread of homospecific Sterile Insect Technique (SIT) as the major approach exploiting sterility in pest control during the last decades [14–18]. It consists of mass rearing, sterilization by ionizing radiation, and massive release of conspecific individuals into the target population. The unfertile mating between the released males and wild females leads to a progressive decline of the target pest population [19,20].

However, a renewed interest has recently been in exploiting reproductive interference to control pest species [8,21–24]. Mitchell et al. [8] reviewed the literature on reproductive interference in natural populations. They highlighted the effects of these processes on population decline in nature and proposed a framework for their use in pest control [8].

Interestingly, McInnis [21] tested the potential use of sterilized males of the oriental fruit fly *Bactrocera dorsalis* (Hendel) against wild carambola fruit fly *B. carambolae*, suggesting that the concern about using non-native or potential pest species to control another pest can be overcome if the released heterospecific individuals are sterile, thus integrating reproductive interference and SIT. More recently, Honma et al. [22] have proposed a framework for incorporating reproductive interference into a classic SIT program. They argued that the sterile males released in a SIT program to suppress the wild-type population of the same species could also lead to suppression of a closely related pest species through reproductive interference (an approach that they called sterile interference).

In this paper, we aimed to explore the potential use of reproductive interference to develop a heterospecific SIT (h-SIT) approach against the spotted wing fly *Drosophila suzukii* Matsumura. This approach, contrary to the classic SIT, is based on using sterile heterospecific males from closely related species to compete with the pest population for mates. *D. suzukii* is an invasive pest that has spread in the last few decades from its native range in East Asia throughout North America, Europe, and South America [25,26]. Unlike most Drosophilidae, *D. suzukii* can lay eggs in unripe and healthy fruits, causing severe economic losses for fruit industries worldwide [27,28]. We selected the fruit fly *D. melanogaster* as the species inducing reproductive interference. Previous studies showed that post-mating isolation between *D. melanogaster* and *D. suzukii* is complete, while there is an incomplete pre-mating isolation [29,24]. Wolf and colleagues, have recently assessed the potential for hybridization between gene drive-modified *D. suzukii* individuals and non-target *Drosophila* species in Europe. They found by male mating behavior tests that *D. melanogaster* males frequently showed interest in *D. suzukii* females but did not achieve copulation [29]. Accordingly, in our previous study, by investigating reproductive interference between non-irradiated *D. melanogaster* males and *D. suzukii*, we found that *D. melanogaster* males successfully courted *D. suzukii* females. Furthermore, they can inseminate *D. suzukii*, leading to egg deposition; however, no hybrids are produced as these eggs did not progress to larval development. Finally, the presence of non-irradiated *D. melanogaster* males under different species ratio also imposed fitness costs on *D. suzukii* females, resulting in reduced *D. suzukii* offspring production [24]. These results are a baseline for exploiting irradiated *D. melanogaster* males in a h-SIT approach.

Here, we specifically aimed: *i*) to assess the effect of irradiation doses on the sterility degree and longevity of *D. melanogaster* males. To this end, we irradiated virgin *D. melanogaster* males with two gamma-ray doses (60 and 80 Gy) and assessed their fertility, through mating trials with *D. melanogaster* females, and their longevity; *ii*) to analyze the mating performance of irradiated *D. melanogaster* males in courting and mating with *D. suzukii* females; *iii*) to evaluate if irradiated *D. melanogaster* males can reduce *D. suzukii* fitness with whom they mated. To this end, we analyzed the effect of irradiated *D. melanogaster* males on the fertility of *D. suzukii* females using different species ratios.

## 2. Material and methods

### 2.1 Laboratory colonies

*Drosophila suzukii* and *D. melanogaster* individuals from the laboratory rearing facilities of the Sapienza University of Rome were used in this study [24]. Both species were reared on an artificial diet consisting of agar (7 g), table sugar (16 g), precooked ground maize (72 g), mother yeast (18 g), soy flour (10 g), and methylparaben (2.5 g) [30]. The colonies were maintained in entomological cages (30 x 30 x 30 cm), in a climate chamber at 25 ±1°C, 14:10 h light:dark cycle.

### 2.2 Effect of irradiation dose on sterility and longevity of *D. melanogaster* males

To evaluate the effect of irradiation doses on the sterility degree of *D. melanogaster* males, mating trials between irradiated males and fertile *D. melanogaster* females were carried out. Virgin *D. melanogaster* adults were obtained by checking the adult emergence from the pupae confined in rearing falcons (50 ml) every thirty minutes, where they completed their full larval development. As soon as new individuals emerged, we divided them into cages (15 x 15 x 15 cm) according to the gender. This procedure allows us to be sure to use only virgin males and females. After selection, *D. melanogaster* males were sterilized using gamma radiation. Males (48-72-96 hours old) were placed in 50 ml falcon tubes containing wet cotton to avoid dehydration and transported to the Calliope Facility at ENEA Casaccia Research Centre (Rome). The Calliope plant is a pool-type irradiation facility equipped with a ^60^Co (mean energy 1.25 MeV) radio-isotopic source array in a high volume (7.0 m ’ 6.0 m ’ 3.9 m) shielded cell. The irradiation cell can provide different dose rates by placing the samples in specific positions and exposing them for varying periods [31]. We provided irradiation doses of 60 Gy and 80 Gy to *D. melanogaster* males, according to Nelson et al. [32] and Henneberry et al. [33]. The dose rate was 175.03 Gy/h (2.92 Gy/min). After irradiation, five irradiated males were placed in falcon (50 ml) with five non-irradiated *D. melanogaster* females. Each falcon contained food substrate to allow females to lay eggs. We assessed two control treatments: a control treatment called “Home”, where we used non-irradiated *D. melanogaster* males, which have not undergone transport stress but have always been kept in the climate chamber conditions; a control treatment called “Trip”, where we used non-irradiated *D. melanogaster* males and females previously transported to the Calliope facility without receiving any irradiation dose. In this way, we evaluated the impact that the stress due to transport could have on their fertility. We performed twelve to sixteen replicates for each irradiation dose (60 and 80 Gy) and the control treatment (0 Gy). The couples were left together for six days and then removed. New individuals that emerged were removed and counted, and oviposition substrates were checked daily until no newborn individuals were observed. We further evaluated the impact of the highest irradiation dose (80 Gy) on the sterility of *Drosophila melanogaster* females, by carrying out mating trials between irradiated females with fertile *D. melanogaster* males as described above. This information can be useful in developing a h-SIT approach since, in mass-rearing conditions, sexing may not be completely accurate, and females could inadvertently be released into the field.

To evaluate the effect of irradiation doses on male longevity, we compared the average lifespan between irradiated and non-irradiated *D. melanogaster* males. Virgin *D. melanogaster* males were selected and irradiated as described above. A cage (30 x 30 x 30 cm) with twenty males was set up for each dose treatment (60 and 80 Gy). Two more cages were set up as control, one as “Control home” and one as “Control trip”, as in the previous experiment. Mortality was recorded every morning until all flies had died.

### 2.3 Mating performance of irradiated *D. melanogaster* males

All the following assays were carried out using only the *D. melanogaster* males irradiated with an 80 Gy irradiation dose, which led to higher sterility without affecting male longevity (see Results section).

First, the mating performance of *D. melanogaster* males was investigated in no-choice and choice tests. In no-choice tests, we set up four experimental conditions by placing in 15 ml falcons: 1) one no-irradiated *D. melanogaster* male and one *D. melanogaster* female; 2) one irradiated *D. melanogaster* male at 80 Gy and one *D. melanogaster* female; 3) one *D. suzukii* male and one *D. suzukii* female; 4) one irradiated *D. melanogaster* male at 80 Gy and one *D. suzukii* female. In this way, it has been possible to understand the average courtship time of *D. melanogaster* males with conspecific and heterospecific females and investigate if the irradiated *D. melanogaster* males court *D. suzukii* females as much as *D. suzukii* males. In the choice test, we set up an experimental condition by placing in 15 ml falcons one *D. suzukii* female, one *D. suzukii* male, and one irradiated *D. melanogaster* male and analyzing the courtship behavior of *D. melanogaster* males and *D. suzukii* males separately when they co-occurred.

The experiments were carried out with virgin individuals selected as described above. After 5 minutes of acclimatation, we recorded 10 minutes of the individual’s behavior with an Olympus Tough TG-6 camera. The videos were used to analyze the courtship behavior elements of *D. melanogaster* and *D. suzukii* males [34,35] using the BORIS software [36]. Twenty replicates were carried out.

Second, we investigated if irradiated *D. melanogaster* males were able to mate with *D. suzukii* females and fecund them. We set up two experimental conditions: - a control condition where we placed in 50 ml falcons with food substrates one virgin *D. suzukii* female alone, to investigate if females lay unfertilized eggs; - an experimental condition where we placed in 50 ml falcons with food substrates, one virgin *D. suzukii* female and one irradiated *D. melanogaster* male at 80 Gy as described previously. We left the individuals inside the falcons for 6 days, to allow the females to oviposit. After that time, we removed the individuals and, through a stereomicroscope Leica EZ4W at magnification 5x we checked daily the possible occurrence of eggs laid by *D. suzukii* females in food substrates. If present, eggs were photographed with a stereomicroscope digital camera and monitored for eclosion and for eventual larval development. Forty replicates were carried out in both conditions.

### 2.4 Effect of irradiated *D. melanogaster* males on *D. suzukii* fitness

We evaluated whether the reproductive interactions between *D. melanogaster* males and *D. suzukii* individuals result in fitness costs for *D. suzukii* females. At this aim, we set up three experimental setting by placing in entomological cages (15 x 15 x 15 cm): 1) five pairs of no-irradiated virgin *D. suzukii* (selected as described above) without *D. melanogaster* males; 2) 40 irradiated *D. melanogaster* males and five pairs of virgin *D. suzukii* to test an 8:1 over-flooding ratio (sterile:wild males, OFR); 3) 60 irradiated *D. melanogaster* males with five pairs of virgin *D. suzukii* to test an OFR of 12:1 [20]. Then, we compared the number of newborn individuals that emerged from cages with only *D. suzukii* pairs and from cages where *D. suzukii* pairs plus irradiated *D. melanogaster* males co-occurred.

### 2.5 Data analysis

To evaluate the effect of the irradiation doses on the sterility degree of *D. melanogaster* adults, a GLM model (generalized linear model; package MASS) [37] was applied. We used the irradiation dose of *D. melanogaster* adults as a fixed effect (0-60-80 Gy) on the offspring produced by *D. melanogaster* females (the response variable), which was recorded as an event with a continuous distribution. We analyzed the offspring produced with a negative binomial distribution applied to the GLM model. The model family was selected comparing the AIC and BIC estimators and the likelihood ratio test. Multiple Comparisons of Means Tukey Contrasts was performed as a post-hoc test using the *multcomp* package in R [38].

To evaluate the effect of the irradiation doses on the longevity of *D. melanogaster* males, survival distributions of the four *D. melanogaster* groups (‘Control cages’, ‘Control trip’, ‘60 Gy’, ‘80 Gy’) were computed using the Kaplan–Meier method [39]. At this end, the *survival* and *survminer* R packages were used [40]. The differences between survival distributions were estimated using the Log-Rank Test.

To analyze the courtship behavior data, we used a GLM model with a negative binomial distribution, according to model selection estimators, for comparing the courtship time among no-choice conditions. As a post-hoc test, we performed the Tukey test. To compare the time spent by *D. melanogaster* and *D. suzukii* males in courting *D. suzukii* females in the choice test, we used the Wilcoxon Mann–Whitney U test (*dplyr* package).

Regarding the effect of *D. melanogaster on D. suzukii* fitness, a GLMM analysis was performed using the *D. suzukii* offspring as a response variable with a continuous distribution. In the analysis, the number of *D. melanogaster* males (0, 40, or 60 individuals) in each replicate was considered a fixed effect (the explanatory variables). The age of the experimental individuals (48, 72, and 96 hours old) was considered a random effect since they are a sampling of infinite possible combinations, and we were not interested in studying them as such but in identifying whether they constituted a source of significant variability. We applied a negative binomial distribution to the GLMM effect, comparing the models based on the optimal parsimony principle (AIC and BIC estimators) and the likelihood ratio test. We performed the Multiple Comparisons of Means Tukey Contrasts as a post-hoc test.

All analyses were performed using the software R version 3.6.2. [41].

## 3. Results

### 3.1 Effect of irradiation dose on sterility and longevity of *D. melanogaster*

In the mating trials aimed to assess the effect of irradiation on male sterility, the mean number of offspring produced by *D. melanogaster* females was 128.66 (± 10.67) (mean ± SE) in the control “Home” and 149.9 (± 18.07) in the control “trip” condition. In the treatments at 60 and 80 Gy, the number of offspring produced by *D. melanogaster* females was 35.81 (± 4.08) and 29.84 (± 3.62), respectively (Fig. 1). The GLM model showed a significant effect of the irradiation dose on the number of offspring produced by *D. melanogaster* females when the males were irradiated at 60 Gy and 80 Gy (Table 1). The Tukey Multiple Comparison tests showed a significant reduction in *D. melanogaster* offspring produced when *D. melanogaster* males were irradiated at 60 Gy (z = −6.268, p = < 0.001) and 80 Gy (z = - 9.335, p = < 0.001) than in the control “home” condition. There was a significant reduction also with irradiated males at 60 Gy (z = −6.784, p = < 0.001) and 80 Gy (z = - 9.702, p = < 0.001) than in the Control “trip” condition. There were no significant differences between the two control conditions (p > 0.05) (Fig. 1).

**Figure 1.**
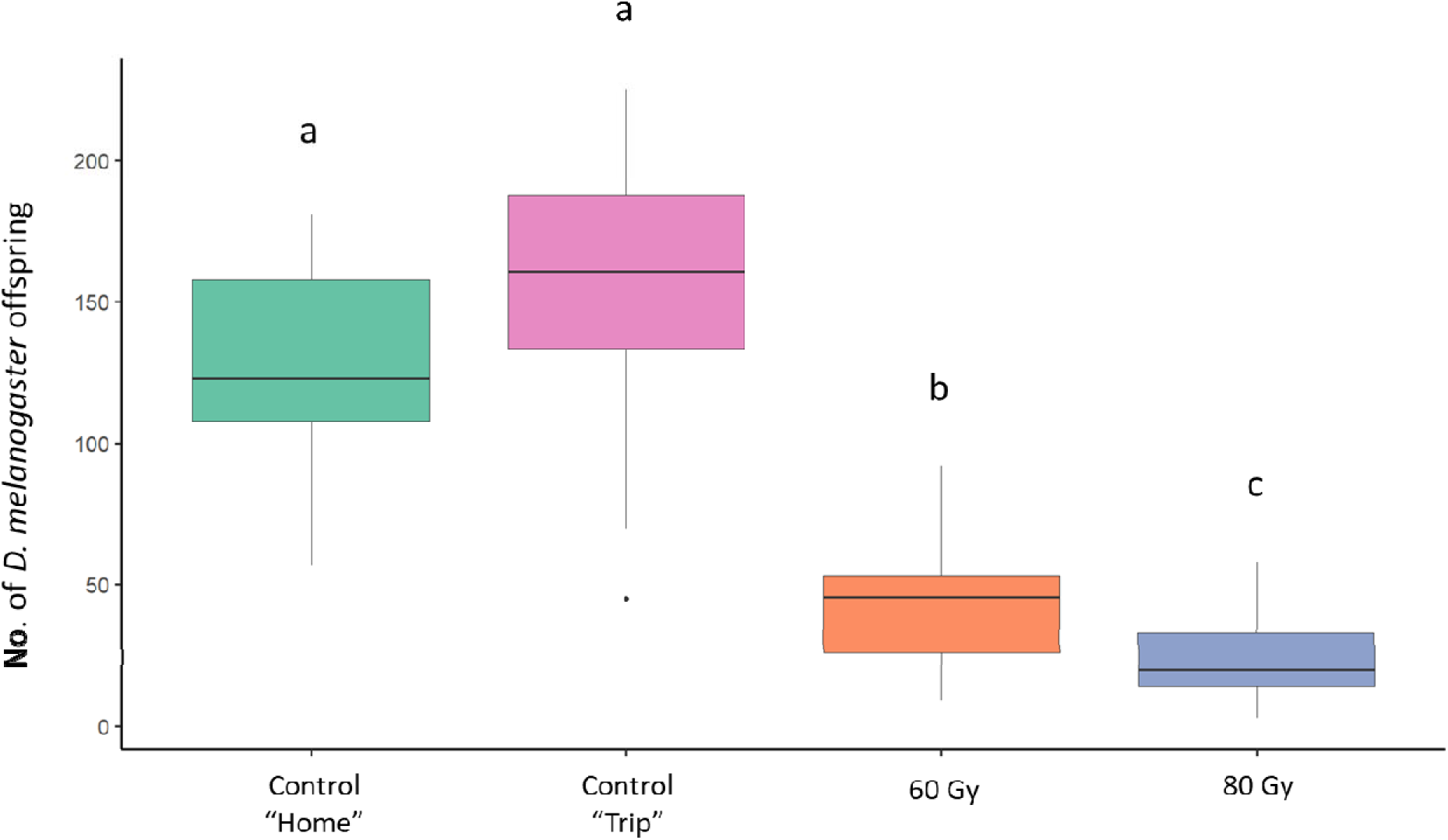
Effect of irradiation on sterility of *Drosophila melanogaster* males. Comparison between the offspring originated from no-irradiated *D. melanogaster* females coupled with no-irradiated *D. melanogaster* males (green column and pink column, respectively) and irradiated at 60 Gy and 80 Gy (orange and blue column, respectively). Different letters mean significant differences by Tukey Multiple Comparison tests (p<0.05).

**Table 1.** Effect of irradiation doses on sterility of *Drosophila melanogaster* males and females. GLM model values are shown. Values in boldface indicate significant differences

| Fixed Effects | Estimate | ±SE | z Value | p-Value |
| --- | --- | --- | --- | --- |
| **Males** |  |  |  |  |
| (Intercept) | 4.8572 | 0.1306 | 37.191 | < 2e-16 |
| Control “Trip” | 0.1527 | 0.1934 | 0.790 | **0.43** |
| IRR60 | -1.1004 | 0.1756 | -6.268 | **3.66e-10** |
| IRR80 | -1.6960 | 0.1817 | -9.335 | < 2e-16 |
| **Females** |  |  |  |  |
| (Intercept) | 4.2850 | 0.1428 | 30.009 | < 2e-16 |
| Control “Trip” | 0.1144 | 0.2016 | 0.568 | **0.57** |
| IRR80 | -1.9741 | 0.2109 | -9.358 | < 2e-16 |

In the mating trials aimed to assess the effect of irradiation on female sterility, the mean number of offspring produced by *D. melanogaster* females was 72.6 (± 10.34) (mean ± SE) in the control “Home” and 81.4 (± 13.86) in the control “trip” condition. In the treatments at 80 Gy, the number of offspring produced by *D. melanogaster* females was 10.08 (± 1.39) (Fig. 2). The GLM model showed a significant effect of the irradiation dose on the number of offspring produced by irradiated *D. melanogaster* females at 80 Gy (Table 1). The Tukey Multiple Comparison tests showed a significant reduction in *D. melanogaster* offspring produced when *D. melanogaster* females were irradiated at 80 Gy (z = - 9.358, p = <1e-04) than in the control “home” condition. There was a significant reduction also with irradiated females at 80 Gy (z = - 9.917, p = <1e-04) than in the Control “trip” condition. There were no significant differences between the two control conditions (p > 0.05) (Fig. 2).

**Figure 2.**
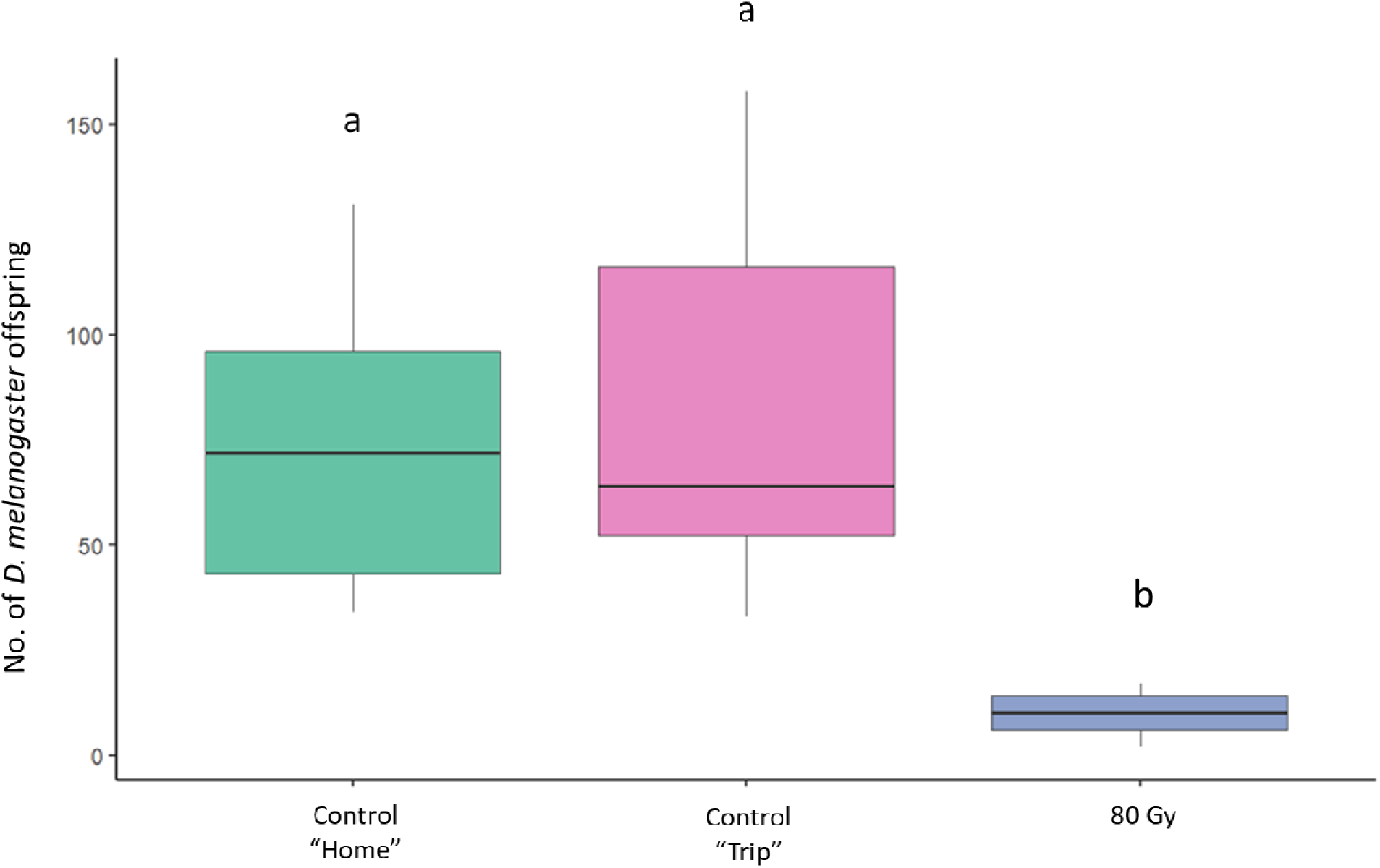
Effect of irradiation on sterility of *Drosophila melanogaster* females. Comparison between the offspring originated from no-irradiated *D. melanogaster* females coupled with no-irradiated *D. melanogaster* males (green and pink columns, respectively) and irradiated female at 80 Gy (blue column). Different letters mean significant differences by Tukey Multiple Comparison tests (p<0.05).

The Kaplan–Meier survival curves showed significant differences in the lifespan of *D. melanogaster* males among treatments (60 Gy, 80 Gy, the Trip cage control and Home cage control groups) (Mantel-Cox log-rank; χ2 = 17.8, d.f.= 3, P = 5e-04) (Fig. 3). The pairwise comparisons test showed that the individuals irradiated with 60 and 80 Gy had higher survival probability than control individuals. Significant differences were indeed observed between the males irradiated at 60 Gy and those of the “Control home” (p= 0.0308) and the “Control trip” (p=0.0151) groups, as well as between the males irradiated at 80 Gy and those of the two control groups (control home p=0.0053; control trip p= 0.0028). No significant differences were observed between the two control groups (p=0.7859) and between the two experimental conditions (p=0.8541).

**Figure 3.**
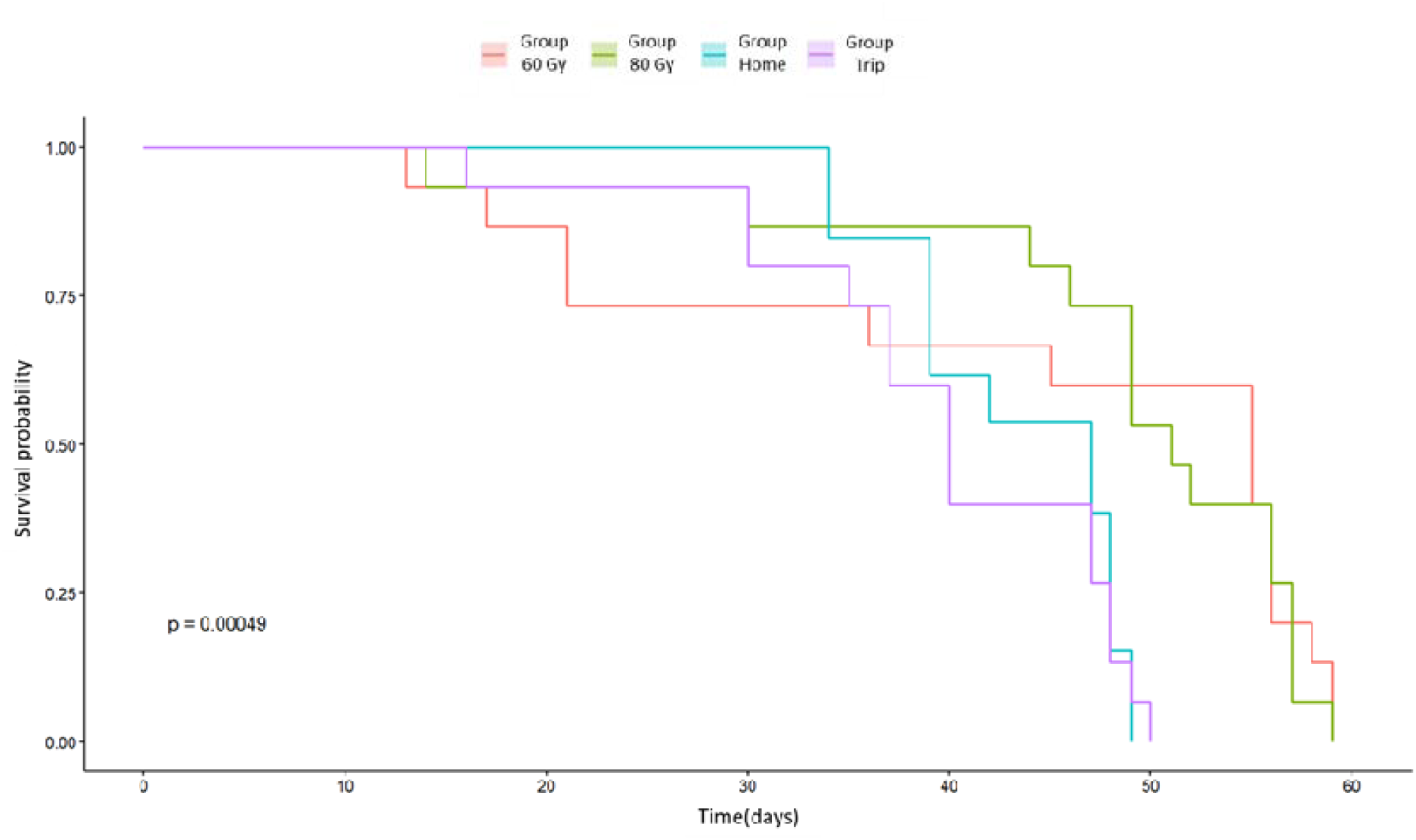
Effect of different treatments on the longevity of *Drosophila melanogaster* males. Kaplan-Meier survival curves for each condition are shown. The p-value of the Log-Rank Test i also shown.

### 3.2 Mating performance of the irradiated *D. melanogaster* males

The mating performance of irradiated *D. melanogaster* males toward *D. suzukii* females was investigated by courtship and mating trials.

First, we investigated if irradiated *D. melanogaster* males were able to court *D. suzukii* females as much as *D. suzukii* males in no-choice and choice tests. Under all experimental conditions the typical behavior elements during courtship were observed, including “orientation” (i.e., the male approaches the female, quivering the abdominal and scissoring its wings), “tapping,” (i.e., the male hits the female abdomen, or middle and hind legs by stretching his foreleg); “wing spreading”, “wing scissoring” (i.e., the male is oriented toward the female front, quivers with the abdomen and scissors his wings keeping them at 180° for seconds to expose the upper side and wing spot toward to female) [34,35].

In the no-choice tests, the average courtship time spent by no-irradiated *D. melanogaster* males courting no-irradiated *D. melanogaster* females was 23.37 % (± 5.19)(mean ± standard error); the average courtship time spent by irradiated *D. melanogaster* males at 80 Gy courting no-irradiated *D. melanogaster* females was 20.84 % (± 3.61); the average courtship time spent by irradiated *D. melanogaster* males courting *D. suzukii* females was 26.41% (± 5.90); the average courtship time spent by *D. suzukii* males courting *D. suzukii* females was 35.36% (± 8.17). The GLM model showed no significant differences in the average courtship time among conditions. The Tukey Multiple Comparison tests showed no significant differences among the four conditions (p> 0.05) (Fig. 4).

**Figure 4.**
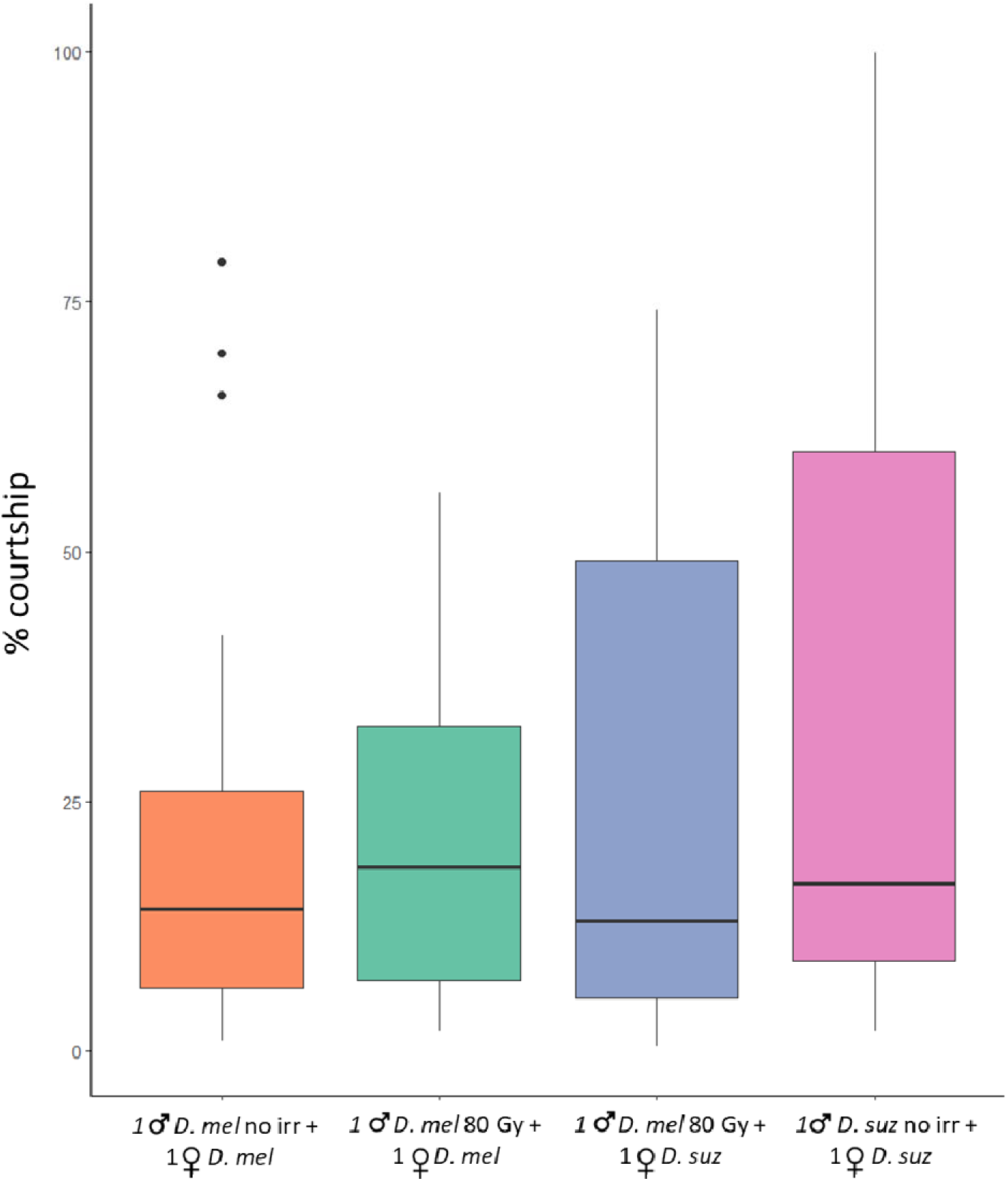
Mating performance of irradiated *Drosophila melanogaster* males in no-choice tests. Time spent in conspecific and heterospecific courtship behavior by no-irradiated and irradiated *D. melanogaster* males and no-irradiated *D. suzukii* males. Percentage of the total time spent courting *D. melanogaster* females by no irradiated *D. melanogaster* males (orange column – conspecific courtship); percentage of the total time spent courting *D. melanogaster* females by irradiated *D. melanogaster* males (green column - conspecific courtship); percentage of the total time spent courting *D. suzukii* females by irradiated *D. melanogaster* males (blue column - heterospecific courtship); percentage of the total time spent courting *D. suzukii* females by no irradiated *D. suzukii* males (pink column - conspecific courtship). Tukey Multiple Comparison tests (p > 0.05). *D. suz* = *D. suzukii*; *D. mel* = *D. melanogaster*.

In choice tests, where *D. melanogaster* males and *D. suzukii* males were placed together in 15 ml falcon with *D. suzukii* females, the average courtship time spent by *D. melanogaster* males was 19.43% (± 6.16) and the average courtship time spent by *D. suzukii* males was 9.52% (± 2.29). There are no significant differences between the courtship time of *D. suzukii* and *D. melanogaster* males (Wilcoxon Mann–Whitney test W = 194.5, p-value = 0.9105) (Fig. 5).

**Figure 5.**
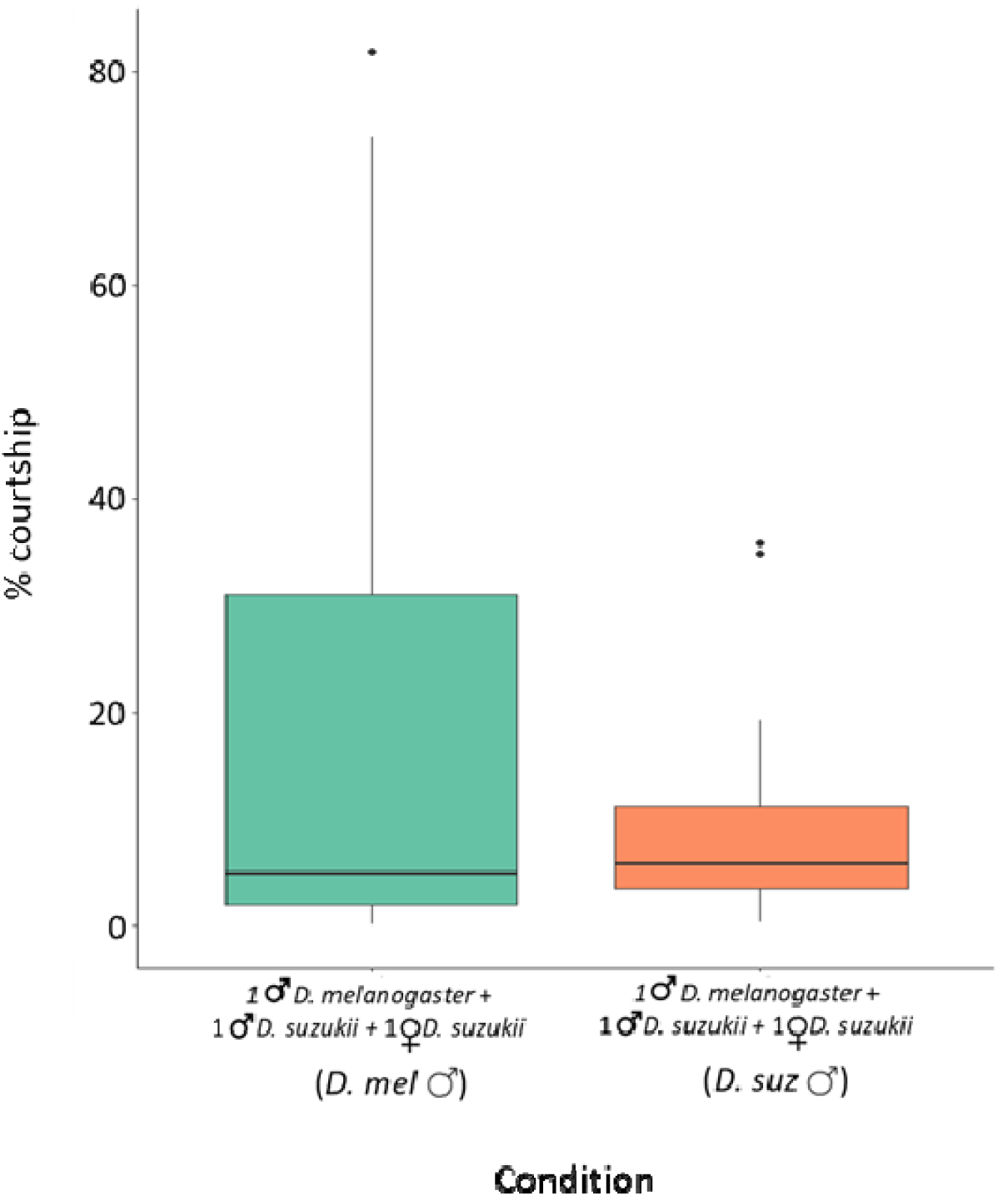
Mating performance of irradiated *Drosophila melanogaster* males in choice tests. Time spent in courting *Drosophila suzukii* females by *D. suzukii* and irradiated *D. melanogaste* males. Percentage of the total time spent courting *D. suzukii* females by irradiated *D. melanogaster* males and *D. suzukii* males when placed together (green and orange column, respectively) Wilcoxon Mann–Whitney test p-value > 0.05.

Second, we investigated if irradiated *D. melanogaster* males could mate and fecund *D. suzukii* females. At this end, we analyzed the occurrence of oviposited eggs by vergin *D. suzukii* females confined alone and with irradiated *D. melanogaster* males. In monitoring the food substrates of each replicate daily, we found no eggs in 39 out of 40 substrates in the condition with only virgin *D. suzukii* females. In the condition with virgin *D. suzukii* females paired with irradiated *D. melanogaster* males, eggs were found in 12 out of 40 replicates (30%), but no subsequent larval development was observed.

### 3.3 Effect of irradiated *D. melanogaster* males on *D. suzukii* fitness

The mean number of the individuals originating from five pairs of *D. suzukii* was 37.95 (± 3.62) (± standard error) in the control tests, while it was 16.11 (± 5.58) and 13.89 (±3.69), when 40 and 60 irradiated *D. melanogaster* males were added, respectively (Fig. 6). The GLMM model showed a significant effect on the *D. suzukii* offspring (Table 2). The Tukey Multiple Comparison tests showed a significant reduction of *D. suzukii* offspring when 40 (z = −2.758, p = 0.01601) and 60 (z = −3.218, p = 0.00376) irradiated *D. melanogaster* males were placed with *D. suzukii* couples (Fig. 6), while no significant differences were observed between treatments with 40 and 60 *D. melanogaster* irradiated males (z = −0.408, p = 0.91176).

**Figure 6.**
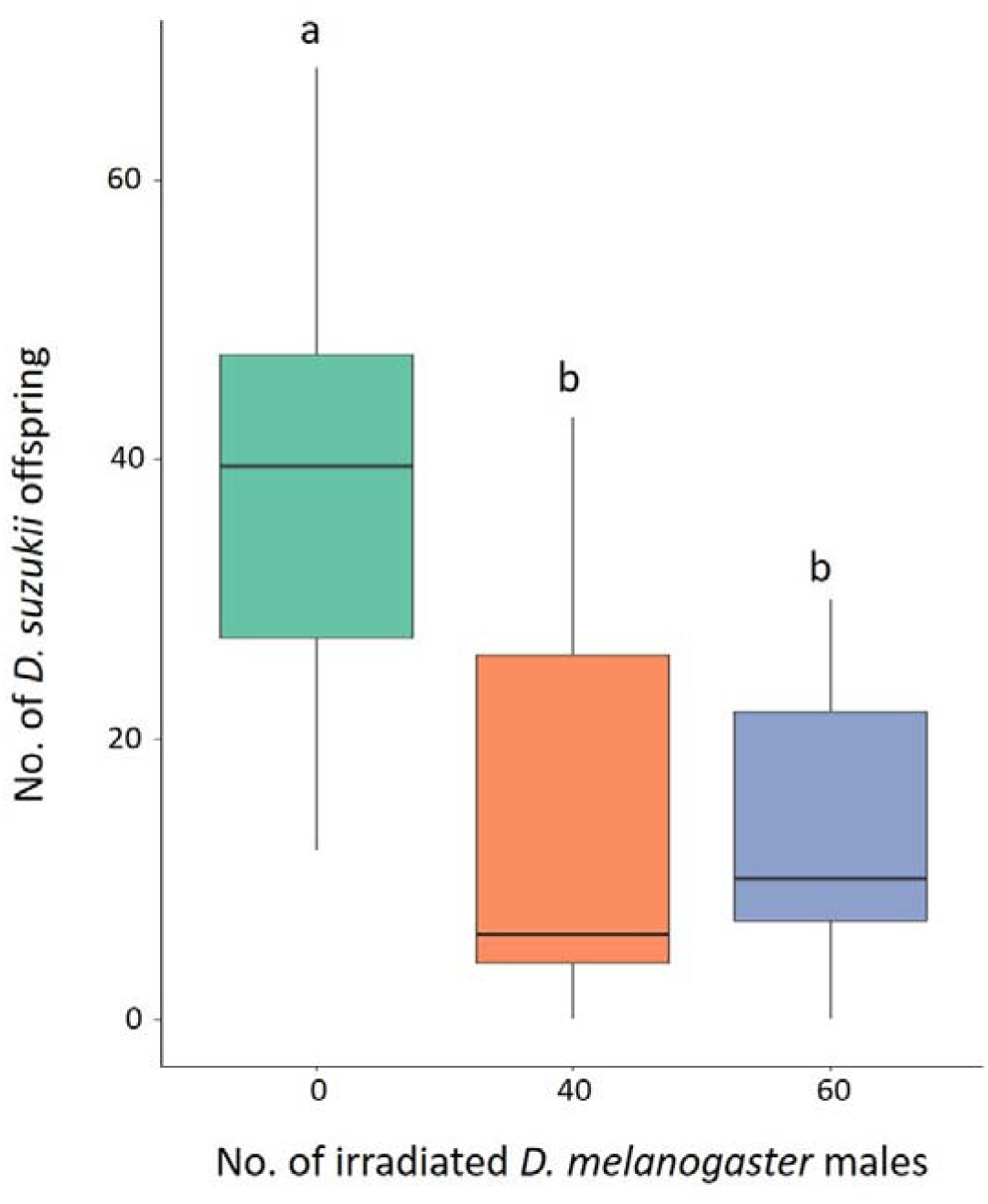
Effect of irradiated *Drosophila melanogaster* males on *D. suzukii* fitness. Offspring originated from five pairs of *D. suzukii* without irradiated *D. melanogaster* males (green column) and with 40 (orange column) or 60 (blue column) males. Different letters mean significant differences by Tukey Multiple Comparison tests.

**Table 2.**
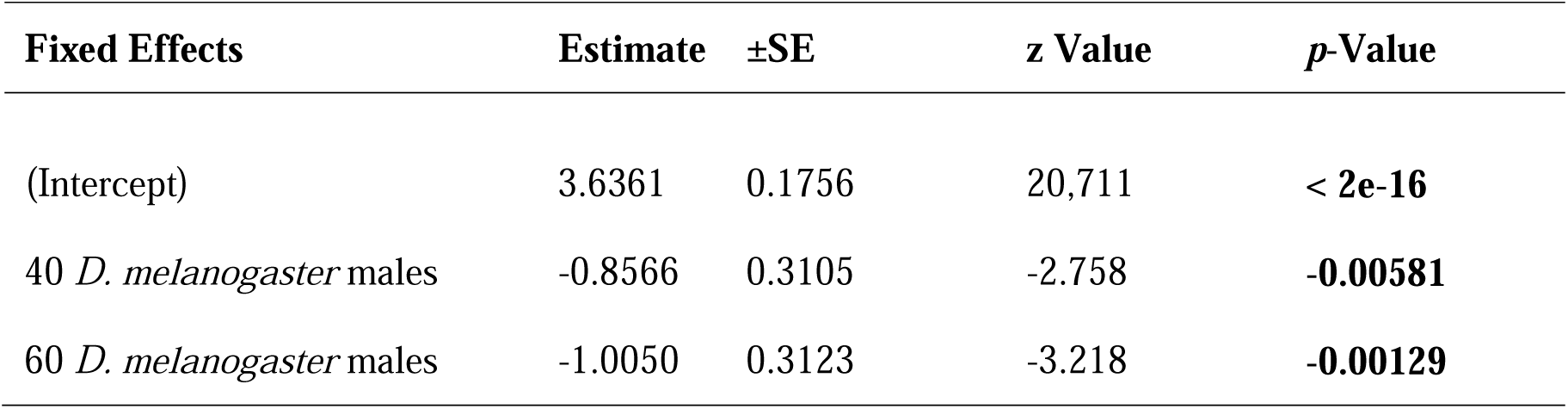
Effect of irradiated *Drosophila melanogaster* males on *D. suzukii* fitness. GLMM model values are shown. Values in boldface indicate significant differences

| Fixed Effects | Estimate | ±SE | z Value | p-Value |
| --- | --- | --- | --- | --- |
| (Intercept) | 3.6361 | 0.1756 | 20,711 | < 2e-16 |
| 40 *D. melanogaster* males | -0.8566 | 0.3105 | -2.758 | **-0.00581** |
| 60 *D. melanogaster* males | -1.0050 | 0.3123 | -3.218 | **-0.00129** |

## 4. Discussion

### 4.1 Irradiation dose, *D. melanogaster* male sterility and longevity

Heterospecific SIT and SIT approaches can be developed under similar theoretical frameworks. Irradiation dose is an important factor affecting the sterility degree, male longevity, and performance [42]. It is, therefore, necessary to find a high enough dose to induce sterility but that has a minimum impact on the biological quality of the irradiated males [43].

We found high levels of sterility in both *D. melanogaster* males and females (Fig. 1 and 2). As observed also in other insect species [44–46], we found that females showed higher sensibility to irradiation than males. While affecting fertility, irradiation did not reduce male longevity. Interestingly, irradiated males exhibited a longer lifespan compared to control individuals. This finding contrasts with some results reported in the literature on Diptera. For example, Lanouette et al. [47] observed no significant differences in the longevity of *D. suzukii* males exposed to increasing irradiation doses (30–120 Gy). In contrast, Chen et al. [48] reported that longevity in *D. suzukii* began to decrease at doses exceeding 90 Gy. Similarly, in *Aedes aegypti*, survival of irradiated males was significantly reduced compared to controls [41,49]. Despite the disparity between our findings and those of other studies, the phenomenon of increased longevity following irradiation has been documented in other species, including some *Anopheles* species [50,51] and the stink bug pest *Bagrada hilaris* [52], where irradiated males at 80 Gy exhibited a longer average lifespan than control individuals. The reasons for such results could be several, since molecular and cellular changes to biological, physical, and human factors [53,54]. Understanding the mechanisms underlying this result would certainly warrant further in-depth analysis. However, within the context of this study, sterile males with stable or extended longevity are likely to remain in the environment longer, effectively competing with fertile males and contributing to population suppression over a prolonged period.

As observed in previous studies [32,33], we also obtained residual fertility in our results. It will be interesting to test if higher irradiation doses will lead to higher male sterility without reducing male performance. Notably, the purpose of male sterilization is different in SIT and h-SIT approaches. In the former, male sterilization is the way to introduce sterility into the wild pest population. Residual fertility could hamper the SIT effectiveness as wild females could mate with released unsterilized males and produce fertile offspring [20,55,56]. Conversely, in the h-SIT, the introduction of sterility in the target pest population is ensured by the reproductive post-zygotic isolation mechanisms between the released heterospecific males and the wild females of the target species [10,21]. Male sterilization avoids potential adverse environmental effects if non-native or potentially pest species are used in the h-SIT. In the case of h-SIT using *D. melanogaster*, residual fertility would not reduce the released males’ effectiveness because not irradiated males would induce sterility in *D. suzukii* females [24]. Furthermore, there are no concerns about the potential release of fertile *D. melanogaster* males. Indeed, *D. melanogaster* is not considered an agricultural pest as the female oviposits on rotten fruits [57].

### 4.2 Mating performance of the irradiated *D. melanogaster* males and effect on *D. suzukii* fitness

Heterospecific SIT is based on the sterility between heterospecifics. The unfertile mating between the released heterospecific males and wild females leads to a progressive decline of the target pest population [19,20]. Therefore, the potential use of h-SIT as a control method strictly depends on the mating ability of the released irradiated males [3]. Our results supported that the irradiated *D. melanogaster* males at 80 Gy can be effective at different stages of mate acquisition, from courtship to mating. The analysis of the courtship behavior showed that *D. melanogaster* males courted *D. suzukii* females as much as *D. suzukii* males (Fig. 4 and 5) Notably, irradiated *D. melanogaster* males exhibited comparable average courtship time toward *D. suzukii* females and conspecific females (Fig. 4). This behavior may be driven by reproductive interference, primarily through misdirected courtship by *D. melanogaster* males. Male insects, including *D. melanogaster*, often exhibit less selective mate choice due to their lower reproductive investment compared to females [58]. This tendency can lead to heterospecific courtship behaviors. Additionally, the larger body size of *D. suzukii* females, often associated with higher fecundity, could make them appear more attractive to *D. melanogaster* males, as body size is commonly used as a proxy for reproductive potential [59,60].

The results of mating trials showed that in the control condition with virgin *D. suzukii* females alone, there were no oviposit eggs in 39 food substrates except for one; larval development and production of *D. suzukii* individuals were observed in that replicate, suggesting we inadvertently selected a *D. suzukii* female that had likely mated before the experiment. Notably, we found that in the test conditions (one *D. suzukii* female placed with one irradiated *D. melanogaster* male), *D. suzukii* females oviposited eggs in 30% of food substrates, and no larval development was observed in any tests. This result confirms that irradiated *D. melanogaster* males could couple, mate, and fecund *D. suzukii* females, but the post-zygotic isolation between *D. suzukii* and *D. melanogaster* is complete [24].

The results of our study clearly show that irradiated *D. melanogaster* males can lead to reproductive interference in *D. suzukii*. Indeed, *D. suzukii* had significantly reduced offspring in the presence of irradiated *D. melanogaster* males, irrespective of the species ratio used (Fig. 6). The reproductive interference on *D. suzukii* by irradiated *D. melanogaster* males is likely due to interference during courtship and mating, as described above. However, other factors could contribute to the offspring reduction of *D. suzukii* by *D. melanogaster* males. First, it has been shown that *D. melanogaster*, during courtship, produces cis-vaccenyl acetate (cVA), which acts as a repellent to *D. suzukii* female for laying eggs [61,62]. Furthermore, *D. melanogaster* males release substances through seminal fluid that reduce female remating in homospecific matings [63,64]. This latter phenomenon deserves future studies. Indeed, if it also occurs in heterospecific matings between *D. suzukii* and *D. melanogaster*, it would be of great interest for heterospecific SIT, preventing *D. suzukii* females from remating with co-specific wild males.

### 4.3 “The importance of being *melanogaster*”

The above arguments make us optimistic about using *D. melanogaster* in h-SIT against *D. suzukii* and fuel to test this approach under well-isolated confined-field facilities.

However, let us highlight a further issue that makes h-SIT particularly exciting to explore: *D. melanogaster* is a model species. The huge amount of data available on the genetics and biology of *D. melanogaster* can be exploited to address classical problems in developing SIT [65,66,67]. Despite considerable research, for most species where SIT is used, we lack a deep understanding of the effects of radiation, rearing conditions (diet, light, density), as well as the effects of environmental conditions on individual biology and male performance in the field. Furthermore, some authors emphasized the need to build “a better male” [68] to overcome the prevailing approach based on overflooding ratios (sterile: wild males) in the field, which can hamper the economic feasibility of the SIT. In this context, *D. melanogaster*, whose biology has been dissected at the molecular, cellular, and physiological levels, could significantly contribute to unveiling these critical issues, making, in perspective, *D. melanogaster*-*suzukii* a model system for the development of heterospecific SIT.

## 5. Conclusions

Reproductive interaction between heterospecific individuals can have multiple effects on pest management. When pre-mating barriers between interfertile species are weak, hybridization can occur and significantly threaten pest control. It can favor pest outbreaks by improving population fitness and adaptation to environmental conditions or by leading to control failures as a consequence of the introgression of pesticide resistance alleles from one species to another [69,70]. On the other hand, reproductive interaction can be beneficial for pest management [8,22]. Here, we highlight the potential use of reproductive interference in pest management by heterospecific SIT. This approach would broaden our control toolbox by offering additional options to control important agricultural pests. In the last decades, the new sequencing technologies combined with classical reproductive incompatibility studies have improved our knowledge of pre- and post-zygotic isolation in sibling pest species, allowing us to identify even more numerous potential systems for the development and application of heterospecific SIT control approach [71,72].

## Acknowledgments

We thank Elisa Michelangeli and Mark Eltenton for their technical help. This study was carried out within the Agritech National Research Center and received funding from the European Union Next-Generation EU (PIANO NAZIONALE DI RIPRESA E RESILIENZA (PNRR) – MISSIONE 4 COMPONENTE 2, INVESTIMENTO 1.4 – D.D. 1032 17 June 2022, CN00000022). This manuscript reflects only the authors’ views and opinions, neither the European Union nor the European Commission can be considered responsible for them. FC was funded by PhD resources within PON “RICERCA E INNOVAZIONE” 2014–2020”, AZIONE IV.5 “DOTTORATI SU TEMATICHE GREEN, D.M. 1061 10 August 2021.

## Data availability

Raw data will be permanently archived if the paper is accepted for publication.

## Author contributions

DP and MC conceived the study. DP, MC and DC designed the study. FC, VM, JS and AV collected the data. FC and VM analyzed the data. DP wrote the first draft of the manuscript. All authors contributed critically to the drafts and approved the final manuscript.

